# Temporal High-Resolution Atlas of Human Blood Leukocyte Composition in Response to Respiratory Virus Inoculation

**DOI:** 10.1101/2024.10.01.616122

**Authors:** Yu Zhang, Tianyi Liu, Chuwen Liu, Xuejun Sun, Pei Liu, James S. Hagood, Raymond J. Pickles, Fei Zou, Xiaojing Zheng

**Affiliations:** Department of Biostatistics, Gillings School of Global Public Health, University of North Carolina at Chapel Hill, Chapel Hill, NC, USA; Department of Probability and Statistics, School of Mathematical Sciences, University of Science and Technology of China, Hefei, Anhui, China; Department of Pediatrics, Division of Pulmonology, School of Medicine, University of North Carolina at Chapel Hill, Chapel Hill, NC, USA; Marsico Lung Institute, School of Medicine, University of North Carolina at Chapel Hill, Chapel Hill, NC, USA; Department of Microbiology & Immunology, School of Medicine, University of North Carolina at Chapel Hill, Chapel Hill, NC, USA; Department of Genetics, School of Medicine, University of North Carolina at Chapel Hill, Chapel Hill, NC, USA; Department of Pediatrics, School of Medicine, University of North Carolina at Chapel Hill, Chapel Hill, NC, USA

## Abstract

**Background:** Respiratory viral infections produce highly variable clinical outcomes, even after controlled exposure. Human viral challenge studies are well suited to study this variation because they capture immune profiles before inoculation and across defined post-challenge intervals. However, how circulating immune-cell populations change over time, and which responses are shared or virus-specific across respiratory viruses, remains incompletely defined.

**Methods:** We leveraged HR-VILAGE-3K3M, a curated human respiratory viral challenge transcriptomic resource, to analyze longitudinal whole-blood profiles from H1N1, H3N2, RSV, and HRV challenge studies. Using statistical deconvolution and xCell-based digital sorting, we estimated changes in major leukocyte populations and 28 immune-cell subtypes in symptomatic infected (Sx/Inf) and asymptomatic uninfected (Asx/Uninf) participants.

Across viruses, symptomatic infection was marked by a broad shared pattern of innate immune activation together with reduced circulating lymphocyte enrichment. Activated dendritic cells, monocytes, and neutrophils increased following challenge, whereas strong pDC activation was observed primarily in influenza studies. Most T-cell and B-cell populations decreased after challenge. Plasma-cell enrichment increased progressively over time. Despite this shared architecture, response timing differed by virus: H3N2 showed the earliest changes, followed by H1N1, whereas RSV responses were delayed and HRV showed more distinct immune-cell trajectories. Migration-associated transcriptional programs were detected across both innate and adaptive immune-cell populations, supporting coordinated immune-cell trafficking during infection. Baseline immune composition also differed between outcome groups, suggesting that pre-existing immune-state differences may influence infection outcomes.

**Conclusions:** These findings define a cellular atlas of human respiratory viral challenge responses and show that respiratory viruses induce a broadly conserved immune-response program with distinct virus-specific kinetics. This framework may help clarify immune features associated with natural resistance, symptomatic infection, and vaccine-relevant antiviral immunity.

## INTRODUCTION

Acute respiratory viral infections remain a major public health burden, yet individuals exposed to the same virus can show markedly different clinical outcomes, ranging from resistance to symptomatic infection. This heterogeneity reflects not only differences in viral factors, but also variation in the host immune state at the time of exposure and in the timing, magnitude, and coordination of subsequent immune responses. Understanding how baseline immune composition and post-exposure immune trajectories differ between protected and susceptible individuals is therefore central to defining mechanisms of natural resistance and symptomatic disease.

Human viral challenge studies provide a uniquely controlled framework for studying these processes. Unlike natural infection cohorts, challenge studies capture immune profiles before inoculation and across well-defined post-exposure intervals, enabling direct comparison of individuals who become symptomatic and infected with those who remain asymptomatic and uninfected. Prior transcriptomic analyses of human challenge cohorts have identified interferon-stimulated, inflammatory, and antiviral gene-expression programs that distinguish symptomatic infection and can predict infection before peak symptoms (Muller et al., 2017; Tsalik et al., 2021; Woods et al., 2013). However, most prior studies have focused primarily on bulk transcriptional signatures or pathway-level responses within individual cohorts, leaving unresolved whether distinct respiratory viruses induce conserved or virus-specific immune-cell trajectories across multiple human challenge studies (Davenport et al., 2015; Huang et al., 2011; Tsalik et al., 2021; Woods et al., 2013).

This question is particularly important because whole-blood transcriptional changes reflect both altered gene expression within cells and shifts in circulating immune-cell composition. Previous studies have reported increased neutrophil and inflammatory signatures together with decreased lymphocyte-associated signatures following respiratory viral infection, but these analyses generally captured broad immune compartments rather than detailed cellular subpopulations (Cheng et al., 2019; Cunha et al., 2009). Experimental cell sorting can resolve immune-cell heterogeneity, but it is difficult to apply retrospectively to large public transcriptomic repositories.

Digital cell-inference approaches provide an alternative strategy by estimating immune-cell proportions or enrichment scores from bulk transcriptomic data, enabling systematic reanalysis of existing human challenge cohorts at higher cellular resolution (L. Dong et al., 2020; M. Dong et al., 2021; T. Liu et al., 2024).

To address this gap, we leveraged HR-VILAGE-3K3M, a curated resource developed by our team that harmonizes publicly available human respiratory viral immunization and challenge transcriptomic datasets across multiple studies (Sun et al., 2025). From this resource, we analyzed longitudinal whole-blood transcriptomic profiles from controlled human viral challenge cohorts spanning H1N1, H3N2, RSV, and HRV. Using complementary statistical deconvolution and xCell-based digital sorting approaches (Aran et al., 2017), we characterized dynamic changes in major leukocyte populations and finer immune-cell subtypes before and after viral challenge. We further compared symptomatic infected participants with asymptomatic uninfected participants to identify immune features associated with susceptibility, protection, and shared versus virus-specific immune-response programs.

Our analyses define a systems-level cellular atlas of respiratory viral challenge responses. We show that baseline immune composition differs between outcome groups before inoculation, suggesting that pre-existing immune-state architecture may contribute to susceptibility to symptomatic infection. Across viruses, we identify conserved innate activation programs, coordinated reductions in circulating T- and B-cell populations, progressive plasma-cell enrichment, and migration-associated transcriptional programs consistent with active immune trafficking. At the same time, the timing and magnitude of these responses differ by virus, with influenza showing earlier innate activation, RSV showing delayed innate immune kinetics, and HRV displaying more distinct response patterns. Together, these findings provide a high-resolution view of shared and virus-specific immune-cell trajectories that shape human responses to respiratory viral challenges.

## METHODS

### Data Curation and Processing

We analyzed blood transcriptomic datasets from human intranasal viral challenge studies originally deposited in the Gene Expression Omnibus (GEO), including GSE73072, a multi-arm viral challenge cohorts with H1N1, H3N2, HRV, and RSV, (T.-Y. Liu et al., 2016; Woods et al., 2013), GSE61754, an H3N2 challenge study (Davenport et al., 2015), and GSE90732, an H1N1 challenge study (Muller et al., 2017). Harmonized quantile-normalized gene-expression matrices and corresponding participant-level metadata were obtained from the HR-VILAGE-3K3M resource, which curates and integrates publicly available respiratory viral challenge transcriptomic datasets across studies (Sun et al., 2025). To ensure biological comparability, analyses were restricted to datasets generated from whole-blood samples.

Participants were classified into asymptomatic/uninfected (Asx/Uninf) and symptomatic infected (Sx/Inf) groups based on infection and symptom metadata provided in the original studies. Infection status was determined based on viral shedding detected 24 hours following challenge, whereas symptom severity was assessed using self-reported clinical symptom scores.

Participants with discordant phenotypes (infected asymptomatic or uninfected symptomatic) were excluded to focus analyses on biologically concordant outcome groups representing susceptibility versus protection phenotypes.

For all longitudinal analyses, observations were restricted to the first 16 days following challenge to minimize the influence of recovery-phase transcriptional changes and sparse sample availability at later time points. In addition, analyses from the GSE73072 H1N1 DEE3 and H3N2 DEE2 cohorts were further restricted to samples collected within 4.5 days post-challenge because of apparent time-dependent batch effects observed beyond this interval (Supp. Fig. 1).

### Cell-Type Deconvolution by NNLS

We performed statistical deconvolution to estimate the proportions of five major leukocyte populations, including T cells, B cells, natural killer (NK) cells, monocytes, and neutrophils. Reference gene-expression profiles for these cell populations were obtained from the Immune Response In Silico (IRIS) dataset (GSE22886), generated using the Affymetrix Human Genome U133A Array (Abbas et al., 2005). Probe annotations were mapped to gene symbols using the corresponding Affymetrix annotation file, yielding 14,387 annotated genes. Among these, 12,174 genes overlapped with the bulk whole-blood transcriptomic datasets used in this study.

To construct a cell-type-specific signature matrix, genes were ranked according to the ratio between expression in the target cell type and the second-highest expressing cell type across the remaining leukocyte populations. The top 150 most cell-type-specific genes were selected and used as the reference signature matrix for NNLS deconvolution analyses.

NNLS deconvolution was applied to whole-blood transcriptomic data from the H1N1 (DEE3), H3N2 (DEE2), and RSV (DEE1) viral challenge cohorts. Samples from asymptomatic/uninfected (Asx/Uninf) and symptomatic infected (Sx/Inf) participants were analyzed separately across representative post-challenge time points (days 0, 0.9, 1.9, 2.9, and 4.5). All analyses were performed in R version 4.3.2 using the NNLS algorithm implemented in the lsei package (Wang et al., 2020).

### Digital Cell-Type Enrichment Analysis by xCell

To estimate the relative enrichment of immune-cell subpopulations from bulk whole-blood transcriptomic data, we applied xCell v1.1(Aran et al., 2017). xCell uses a gene signature–based framework built upon single-sample gene set enrichment analysis (ssGSEA) (Barbie et al., 2009) and incorporates spillover correction between closely related cell types. Unlike constrained deconvolution approaches, xCell generates independent enrichment scores rather than estimated cell proportions.

Twenty-eight leukocyte subpopulations relevant to peripheral blood were selected from the xCell reference framework for downstream analyses (Supp. Table 1).

### Meta-analysis of Temporal Immune Responses across Studies

To evaluate temporal immune-response dynamics following viral challenge, sampling time points were grouped into half-day intervals while retaining pre-challenge day 0 as a shared baseline reference across studies.

Within each study, linear regression models were fit separately for each pathogen, cell type, and time interval:

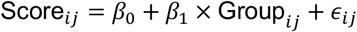

where Score_*ij*_ represents the xCell enrichment score for sample *j* in study *i*, and Group _*ij*_ denotes symptomatic infected (Sx/Inf) versus asymptomatic/uninfected (Asx/Uninf) status. The coefficient *β*_1_ therefore represented the estimated enrichment difference between outcome groups.

When multiple studies contributed to the same pathogen–cell-type–time-point combination, study-specific regression coefficients were combined using inverse-variance-weighted fixed-effect meta-analysis:

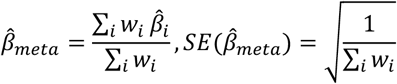

where 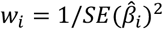 represents the inverse-variance weight for study *i*. This approach generated pooled effect estimates and corresponding 95% confidence intervals for each pathogen, cell type, and time interval.

When only a single study was available for a given pathogen–cell-type–time-point combination, study-specific estimates were retained without pooling. Because RSV included only one eligible study, RSV effect estimates were therefore reported directly without meta-analysis.

### Migration and Residency Score Analysis

To evaluate migration- and residency-associated transcriptional programs, we obtained curated migration-marker genes (e.g., *CX3CR1, KLF2*) and residency-marker genes (e.g., *CD69, CXCR6*) from Zhao et al. (2021). Migration and residency scores were calculated using the AddModuleScore function in Seurat v4.3 (Hao et al., 2021), which computes normalized expression scores for predefined gene sets relative to randomly selected control genes.

Associations between migration or residency scores and xCell-derived immune-cell enrichment scores were evaluated using mixed-effects regression models with first-order autoregressive [AR(1)] covariance structures and restricted maximum likelihood (REML) estimation. Time and cell-type enrichment scores were included as fixed effects, whereas participant-specific random intercepts were included to account for repeated longitudinal measurements. Wald tests were used to assess fixed-effect significance, and multiple-testing correction was performed using the Benjamini–Hochberg false discovery rate (FDR) procedure (Benjamini & Hochberg, 1995).

These analyses were performed using the H1N1 (DEE3), H3N2 (DEE2), and RSV (DEE1) whole-blood viral challenge cohorts, consistent with the datasets used for deconvolution analyses.

## RESULTS

### Data Curation and Study Selection for Viral Challenge Analysis

A total of nine human viral challenge cohorts representing four respiratory viruses, H1N1, H3N2, RSV, and HRV, were included from the large-scale GSE73072 study together with independent H1N1 and H3N2 challenge studies (GSE90732 and GSE61754). All studies included longitudinal whole-blood transcriptomic profiling with pre-challenge baseline and serial post-challenge sampling. Participants were stratified into asymptomatic/uninfected (Asx/Uninf) and symptomatic infected (Sx/Inf) groups based on available infection and symptom outcome metadata. A detailed summary of included studies is provided in Table 1.

**Table 1.** Summary of Challenge Studies and Sample Metadata.

| Viral Challenge | Tissue | Sample Size | Timepoints Used in Analysis (days) | Platform | GEO Accession |
| --- | --- | --- | --- | --- | --- |
| H1N1 | Whole blood | 9 Sx/Inf,<br>6 Asx/Uninf | -0.9, 0.0, 0.2, 0.5, 0.9, 1.2, 1.5, 1.9, 2.2, 2.5, 2.9, 3.2, 3.5, 3.9, 4.2, 4.5 | Affymetrix Human Genome U133A 2.0 Array | GSE73072 DEE3 |
| H1N1 | Whole blood | 6 Sx/Inf<br>3 Asx/Uninf | -0.9, 0.0, 0.2, 0.5, 0.9, 1.2, 1.5, 1.9, 2.2, 2.5, 2.9, 3.2, 3.5, 3.9, 4.2, 4.5, 4.9, 5.2, 5.5, 5.9, 6.9 | Affymetrix Human Genome U133A 2.0 Array | GSE73072 DEE4 |
| H1N1 | Whole blood | 8 Sx/Inf,<br>3 Asx/Uninf | 0.0, 1.0, 2.0, 3.0, 4.0 | Illumina HumanHT-12 V4.0 expression beadchip | GSE90732 |
| H3N2 | Whole blood | 9 Sx/Inf,<br>6 Asx/Uninf | -1.0, 0.0, 0.2, 0.5, 0.9, 1.2, 1.5, 1.9, 2.2, 2.5, 2.9, 3.2, 3.5, 3.9, 4.2, 4.5 | Affymetrix Human Genome U133A 2.0 Array | GSE73072 DEE2 |
| H3N2 | Whole blood | 8 Sx/Inf<br>6 Asx/Uninf | -1.2, 0.1, 0.4, 0.8, 1.1, 1.4, 1.8, 2.1, 2.4, 2.8, 3.1, 3.4, 3.8, 4.1, 4.4, 4.8, 5.1, 5.4, 5.8, 6.1, 6.8, 7.1 | Affymetrix Human Genome U133A 2.0 Array | GSE73072 DEE5 |
| H3N2 | Whole blood | 8 Sx/Inf<br>3 Asx/Uninf | 0.0, 0.5, 1.0, 2.0 | Illumina HumanHT-12 V4.0 expression beadchip | GSE61754 |
| HRV | Whole blood | 12 Sx/Inf<br>4 Asx/Uninf | -0.5, 0.0, 0.2, 0.3, 0.5, 0.7, 0.8, 1.0, 1.2, 1.5, | Affymetrix GeneChip Human Genome | GSE73072 Duke |
|  |  |  | 1.8, 2.0, 2.5, 3.0, 3.5, 4.0, 4.5, 5.0, 5.7 | U133A 2.0 Array |  |
| HRV | Whole blood | 10 Sx/Inf<br>6 Asx/Uninf | -0.5, 0.0, 0.2, 0.3, 0.5, 0.7, 0.8, 1.0, 1.2, 1.5, 1.8, 2.0, 3.0, 4.0, 5.0 | Affymetrix GeneChip Human Genome U133A 2.0 Array | GSE73072<br>UVA |
| RSV | Whole blood | 10 Sx/Inf,<br>7 Asx/Uninf | -1.6, 0.0, 0.2, 0.5, 0.9, 1.2, 1.5, 1.9, 2.2, 2.5, 2.9, 3.2, 3.5, 3.9, 4.2, 4.5, 4.9, 5.2, 5.5, 5.9, 6.9 | Affymetrix GeneChip Human Genome U133A 2.0 Array | GSE73072<br>DEE1 |

### Baseline immune composition differs before challenge and may shape susceptibility

We first used statistical deconvolution to estimate the baseline proportions of five major immune-cell populations (T cells, B cells, NK cells, monocytes, and neutrophils) across H3N2, H1N1, and RSV challenge studies. Prior to challenge, Sx/Inf participants tended to exhibit higher neutrophil and monocyte proportions and lower B-cell and T-cell proportions compared with Asx/Uninf participants, particularly in the H3N2 study, although statistically significant differences were observed only for neutrophils (Fig. 1).

**Figure 1.**
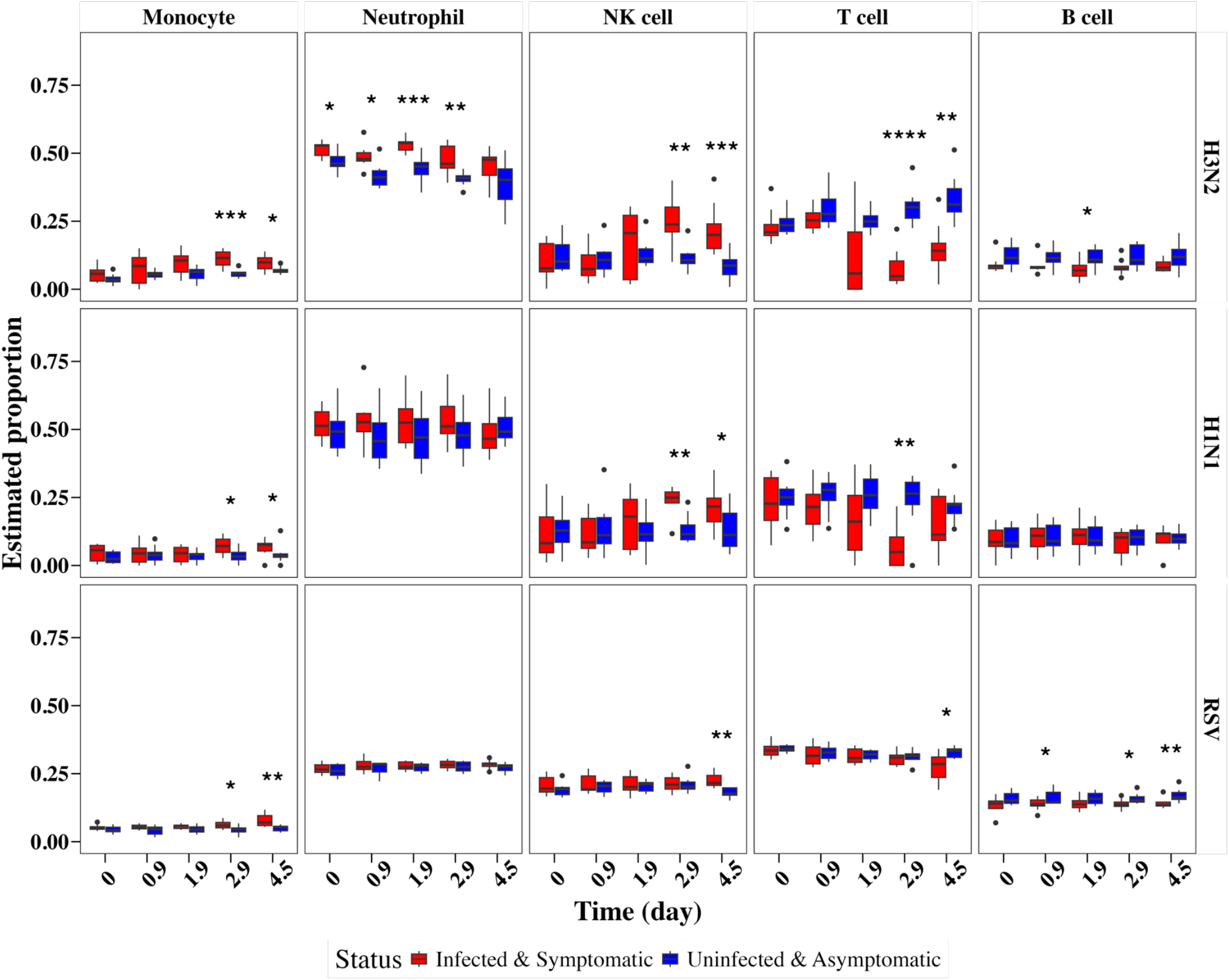
Boxplots showing the deconvoluted cell type proportions of samples from the H3N2 (top), H1N1 (middle), and RVS (bottom) studies at various time points (in days) among infected and symptomatic participants (red) and uninfected and asymptomatic participants (blue). The cell types, from left to right, are monocytes, neutrophils, natural killer (NK) cells, T cells, and B cells. Statistical significance was assessed using the Wilcoxon rank-sum test and is indicated by the following symbols: * (0.01 ≤ p < 0.05), ** (0.001 ≤ p < 0.01), *** (10^-4 ≤ p < 0.001), and **** (p < 10^-4).

To further resolve the immune-cell subpopulations underlying these broad baseline immune differences, we next applied xCell digital cell-type enrichment analysis across 28 blood leukocyte subpopulations. Compared with the major-cell-type deconvolution framework, xCell enabled higher-resolution characterization of baseline immune-state architectures across innate and adaptive immune compartments.

Consistent with the deconvolution findings, several lymphocyte subpopulations inferred by xCell, including CD8^+^ T cells and B cells, showed lower baseline enrichment in Sx/Inf participants in selected viral studies, particularly in H3N2 and RSV cohorts. In contrast, several innate immune-cell populations, including neutrophils, tended to exhibit higher baseline enrichment among Sx/Inf participants. Together, these findings suggest that baseline immune-state are different between Sx/Inf and Asx/Uninf participants prior to challenge.

## Conserved cross-virus innate immune activation programs with distinct temporal kinetics

To characterize longitudinal innate immune dynamics following respiratory viral challenge, we analyzed xCell-derived enrichment trajectories across 28 leukocyte subpopulations in H1N1, H3N2, HRV, and RSV challenge studies (Figs. 2–3). When multiple studies were available for the same pathogen–cell type–time-point combination, estimates were combined using fixed-effect meta-analysis.

**Figure 2.**
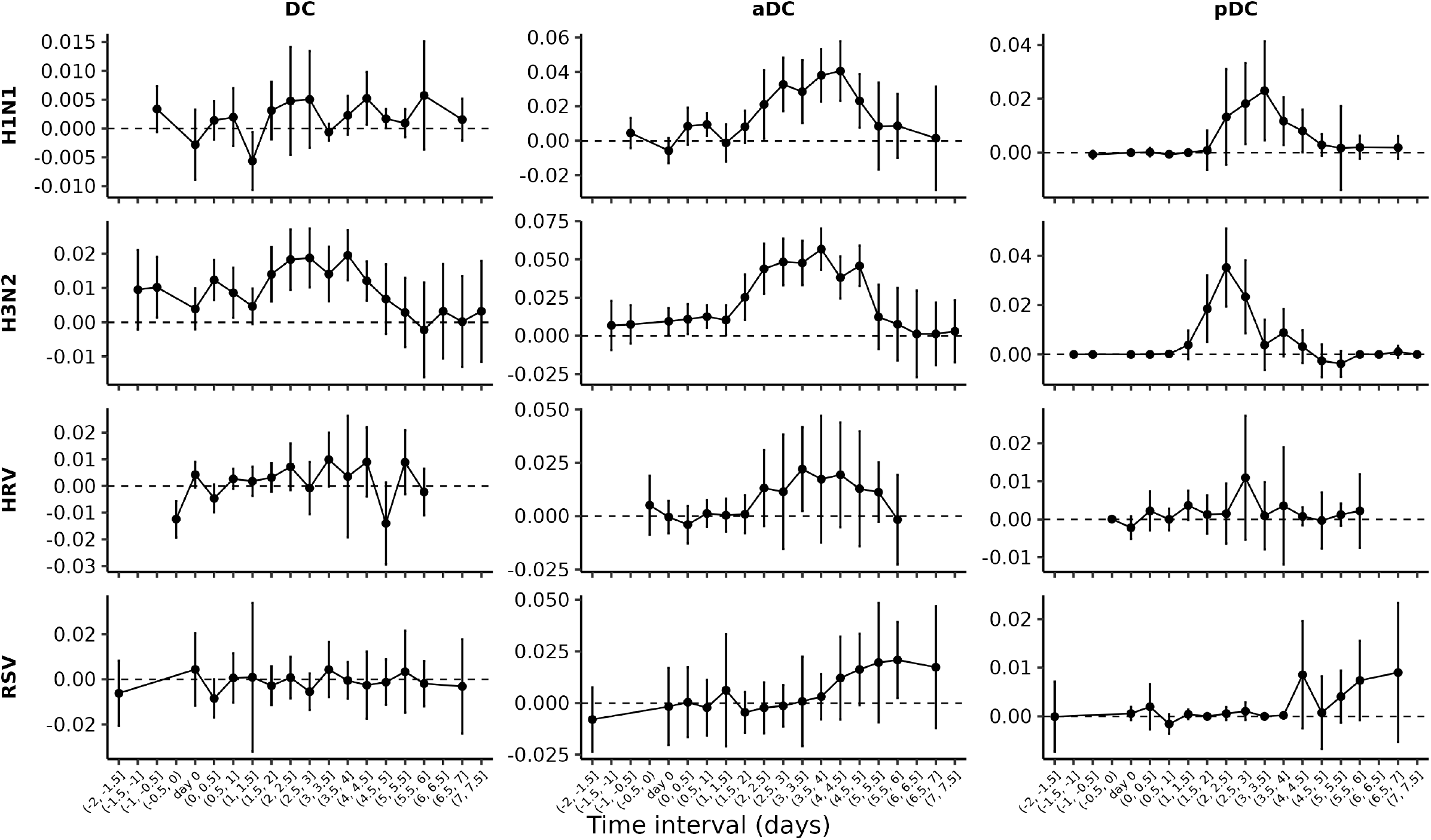
Fixed-effect meta-analysis of dendritic cell subtype enrichment following viral inoculation. Columns represent dendritic cell subtypes— conventional dendritic cells (DC, left), activated dendritic cells (aDC, middle), and plasmacytoid dendritic cells (pDC, right). Rows correspond to viral pathogens, including H1N1, H3N2, HRV, and RSV (top to bottom). Each point shows the pooled effect size estimate from the fixed-effect meta-analysis comparing symptomatic/infected (Sx/Inf) versus asymptomatic/uninfected (Asx/Uninf) participants at each 0.5-day time interval. Vertical error bars denote 95% confidence intervals (CIs); intervals that do not cross zero indicate statistically significant differences between the two groups. A positive estimated value reflects higher relative enrichment in the Sx/Inf group. Day 0 represents the baseline time point prior to or at inoculation.

**Figure 3.**
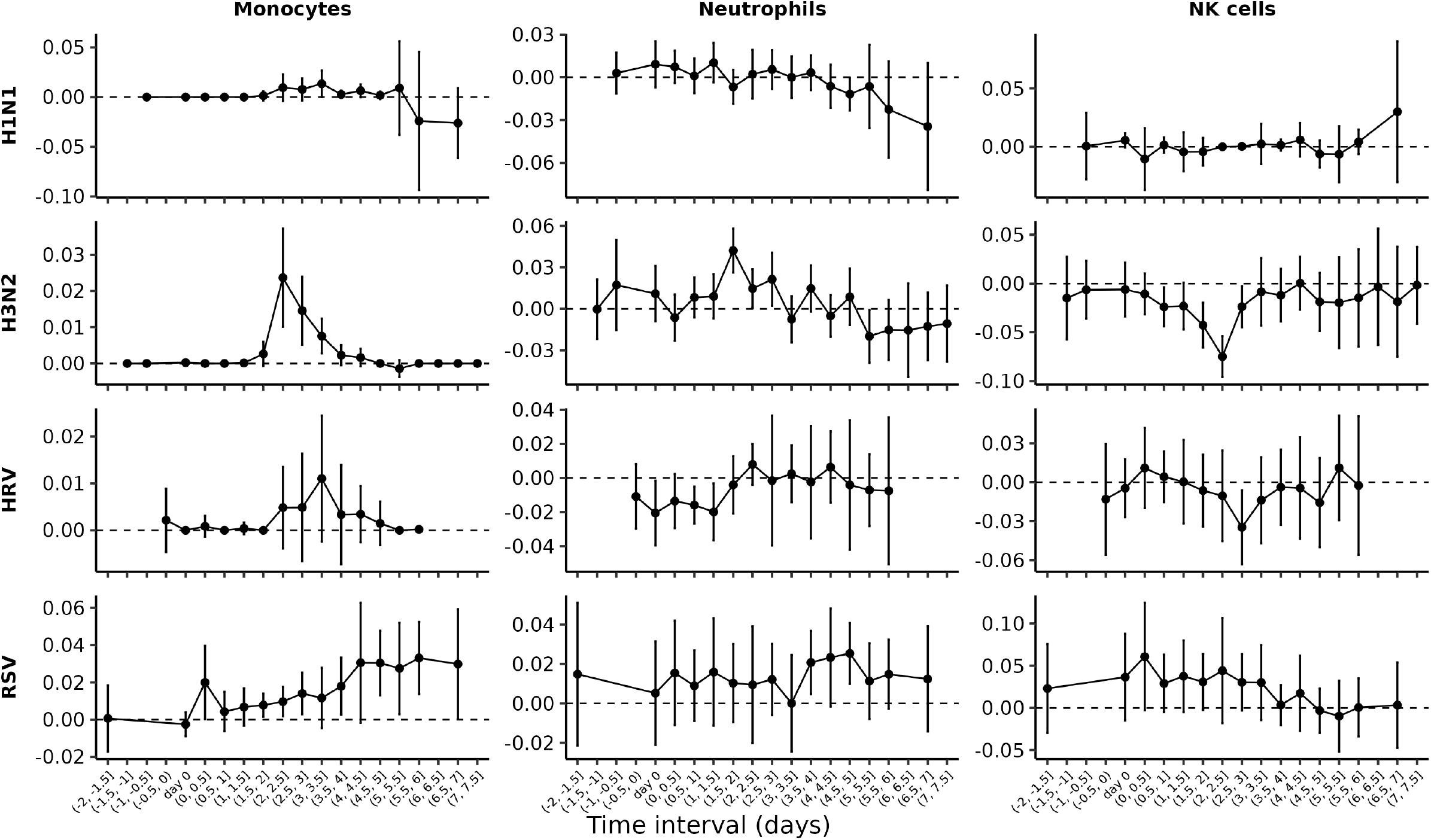
Fixed-effect meta-analysis of dendritic cell subtype enrichment following viral inoculation. Columns represent monocytes (left), neutrophils (middle), and natural killer (NK) cells (right). Rows correspond to viral pathogens, including H1N1, H3N2, HRV, and RSV (top to bottom). Each point shows the pooled effect size estimate from the fixed-effect meta-analysis comparing symptomatic/infected (Sx/Inf) versus asymptomatic/uninfected (Asx/Uninf) participants at each 0.5-day time interval. Vertical error bars denote 95% confidence intervals (CIs); intervals that do not cross zero indicate statistically significant differences between the two groups. A positive estimated value reflects higher relative enrichment in the Sx/Inf group. Day 0 represents the baseline time point prior to or at inoculation.

Among innate immune-cell populations, activated dendritic cells (aDCs) showed some of the earliest and strongest temporal responses. Significant increases in aDC enrichment among Sx/Inf comparing to Asx/Uninf were detected as early as days 1.5 and 2 following H3N2 and H1N1 challenge respectively, peaking around days 3.5–4. In HRV studies, aDC enrichment increased beginning around day 2.5, although statistically significant differences were not observed until days 3–3.5 post-challenge. RSV demonstrated a similar but delayed pattern, with significant aDC enrichment emerging around day 6 post-challenge.

Plasmacytoid dendritic cells (pDCs) exhibited temporal activation patterns closely paralleling those of aDCs following H1N1 and H3N2 challenge. In contrast, pDC enrichment remained relatively unchanged following HRV and RSV challenge between Sx/Inf and Asx/Uninf. Similarly, dendritic cells (DCs) showed significant increase in enrichment among Sx/Inf approximately 2–4 days following H3N2 challenge, whereas corresponding increases following H1N1 challenge were weaker and not statistically significant.

Monocyte enrichment increased in Sx/Inf participants following H3N2 and RSV challenge but showed weaker or nonsignificant changes in H1N1 and HRV studies. Neutrophils demonstrated rapid early increases following H3N2 challenge, with significant enrichment detected as early as day 1.5, whereas RSV-associated neutrophil enrichment occurred later, around days 4–5 post-challenge. In contrast, HRV studies showed transient decreases in neutrophil enrichment shortly after challenge, whereas no significant neutrophil changes were identified following H1N1 challenge.

NK-cell enrichment decreased significantly in Sx/Inf participants between days 2–4 following H3N2 challenge and around day 3 following HRV challenge, whereas no significant differences were observed in H1N1 or RSV studies.

These findings demonstrate that innate immune-cell populations exhibit broadly conserved infection-induced activation patterns across respiratory viruses, although the temporal kinetics of these responses differ substantially between viruses. The earliest responses were observed following H3N2 challenge, whereas RSV exhibited the most delayed innate immune activation kinetics.

### Coordinated innate-to-adaptive immune transitions across respiratory viruses

We next examined longitudinal changes in lymphocyte subpopulations using xCell digital sorting (Figs. 4–7). Consistent with the deconvolution analyses, most T-cell populations showed decreased enrichment in Sx/Inf participants compared with Asx/Uninf participants following viral challenge.

**Figure 4.**
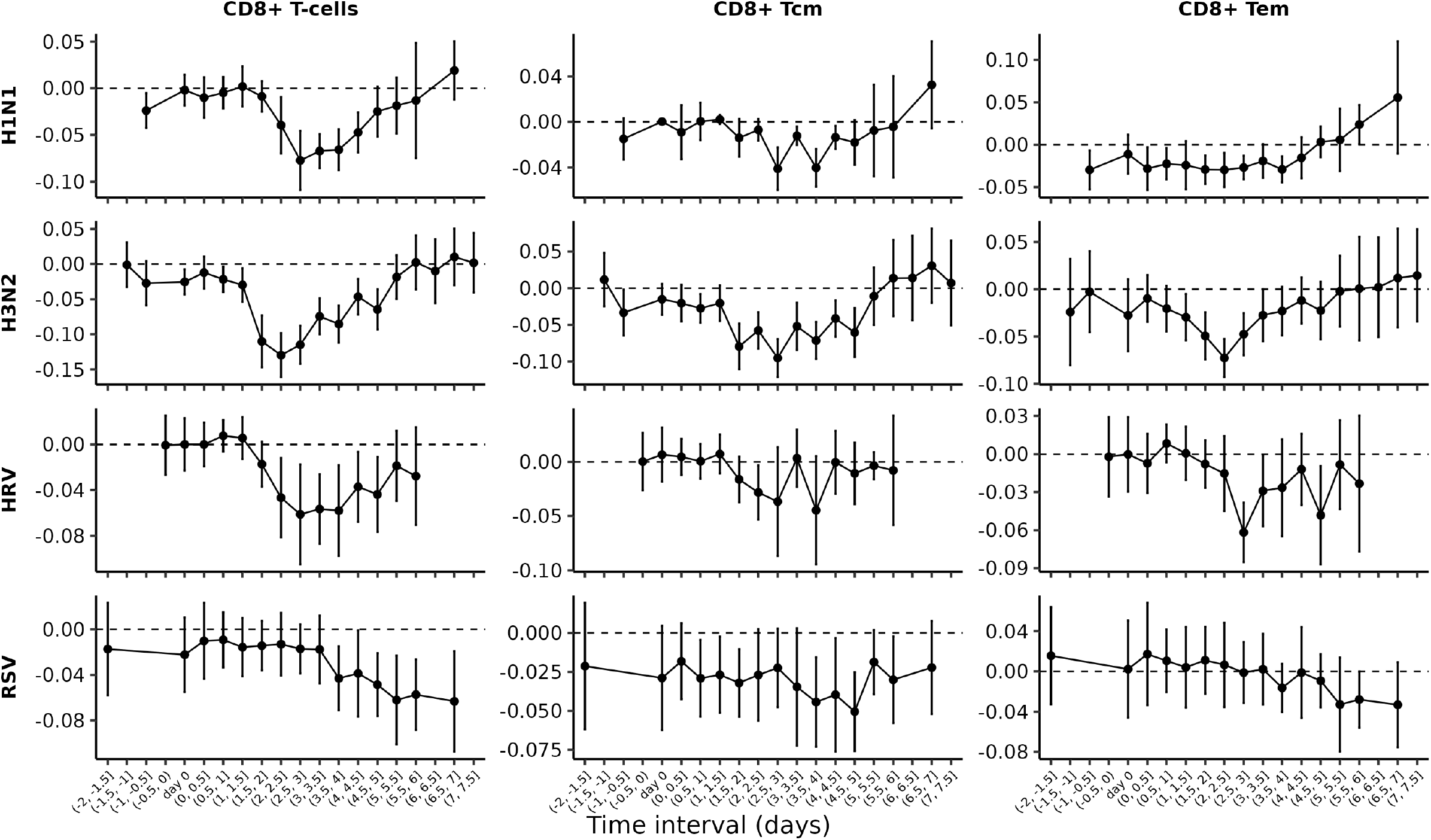
Fixed-effect meta-analysis of dendritic cell subtype enrichment following viral inoculation. Columns represent CD8+ T cells (left), CD8+ central memory T cells (CD8+ Tcm, middle), and CD8+ effector memory T cells (CD8+ Tem, right). Rows correspond to viral pathogens, including H1N1, H3N2, HRV, and RSV (top to bottom). Each point shows the pooled effect size estimate from the fixed-effect meta-analysis comparing symptomatic/infected (Sx/Inf) versus asymptomatic/uninfected (Asx/Uninf) participants at each 0.5-day time interval. Vertical error bars denote 95% confidence intervals (CIs); intervals that do not cross zero indicate statistically significant differences between the two groups. A positive estimated value reflects higher relative enrichment in the Sx/Inf group. Day 0 represents the baseline time point prior to or at inoculation.

CD8^+^ T-cell enrichment decreased earliest following H3N2 challenge, with significant reductions observed around days 1.5–2 post-challenge that persisted through day 5. Similar but later decreases were observed following H1N1 and HRV challenge, whereas RSV responses occurred substantially later. CD8^+^ central memory T cells (CD8^+^ Tcm) showed similar temporal patterns, with reductions beginning around day 2 after H3N2 challenge and around day 3 following H1N1 challenge. In RSV studies, CD8^+^ Tcm enrichment remained consistently lower in Sx/Inf participants before and after challenge, although statistically significant differences were observed mainly at later post-challenge time points.

CD8^+^ effector memory T cells (CD8^+^ Tem) also decreased following H3N2 and HRV challenge. In H1N1 studies, CD8^+^ Tem enrichment remained consistently lower among Sx/Inf participants both before challenge and during the early post-challenge period, whereas RSV studies showed minimal significant changes.

CD4^+^ T-cell populations broadly mirrored the CD8^+^ T-cell trajectories. Significant reductions in CD4^+^ T cells and Th2 cells were observed across all four viral challenge models (Fig. 5). CD4^+^ naïve, central memory, and effector memory T-cell subsets also declined with similar virus-specific temporal patterns, occurring earliest in H3N2 studies and latest in RSV studies. In contrast, Th1-cell enrichment increased around day 4 following H3N2 challenge.

**Figure 5.**
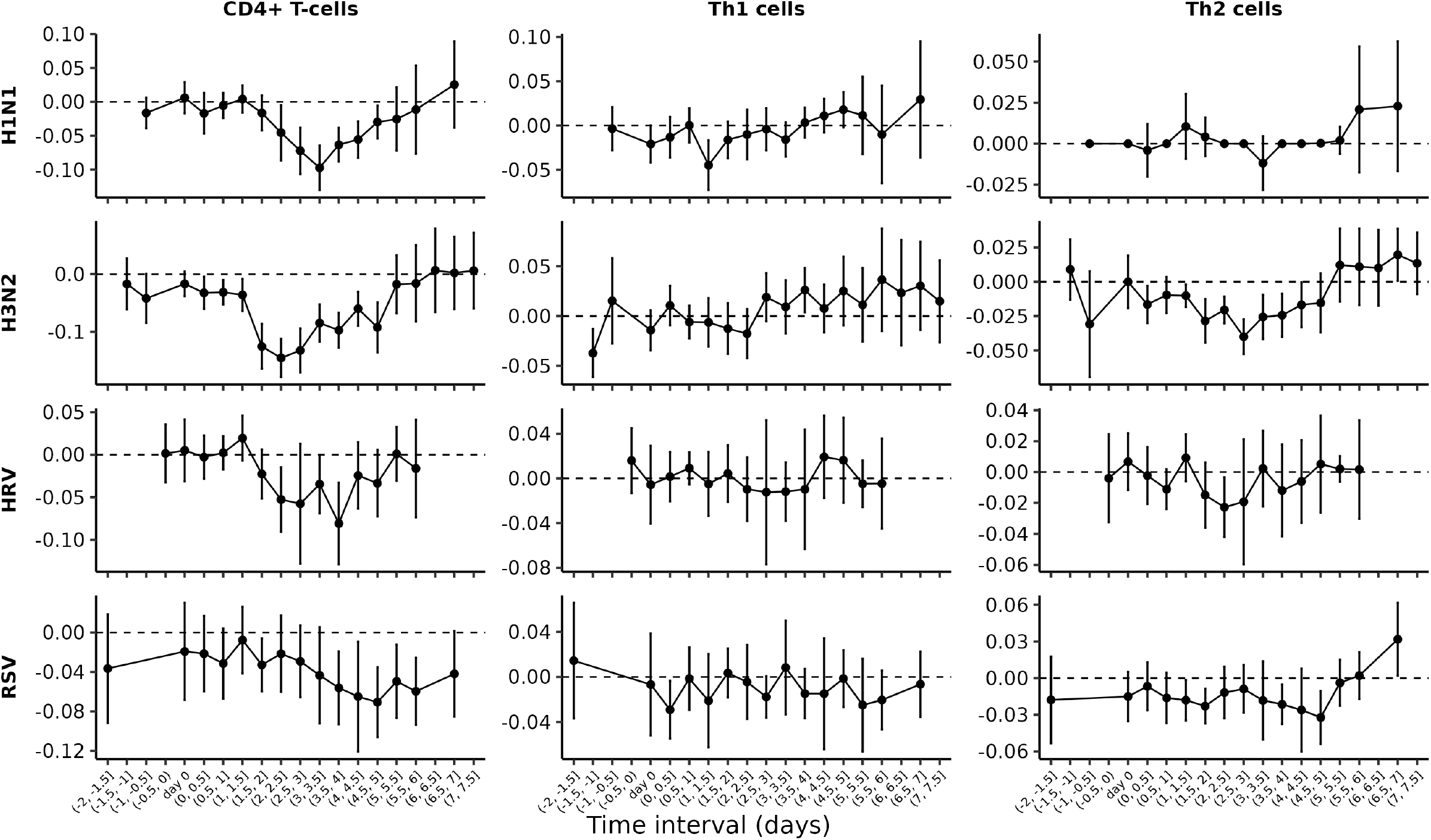
Fixed-effect meta-analysis of dendritic cell subtype enrichment following viral inoculation. Columns represent CD4+ T cells (left), T helper 1 (Th1) cells (middle), and T helper 2 (Th2) cells (right). Rows correspond to viral pathogens, including H1N1, H3N2, HRV, and RSV (top to bottom). Each point shows the pooled effect size estimate from the fixed-effect meta-analysis comparing symptomatic/infected (Sx/Inf) versus asymptomatic/uninfected (Asx/Uninf) participants at each 0.5-day time interval. Vertical error bars denote 95% confidence intervals (CIs); intervals that do not cross zero indicate statistically significant differences between the two groups. A positive estimated value reflects higher relative enrichment in the Sx/Inf group. Day 0 represents the baseline time point prior to or at inoculation.

**Figure 6.**
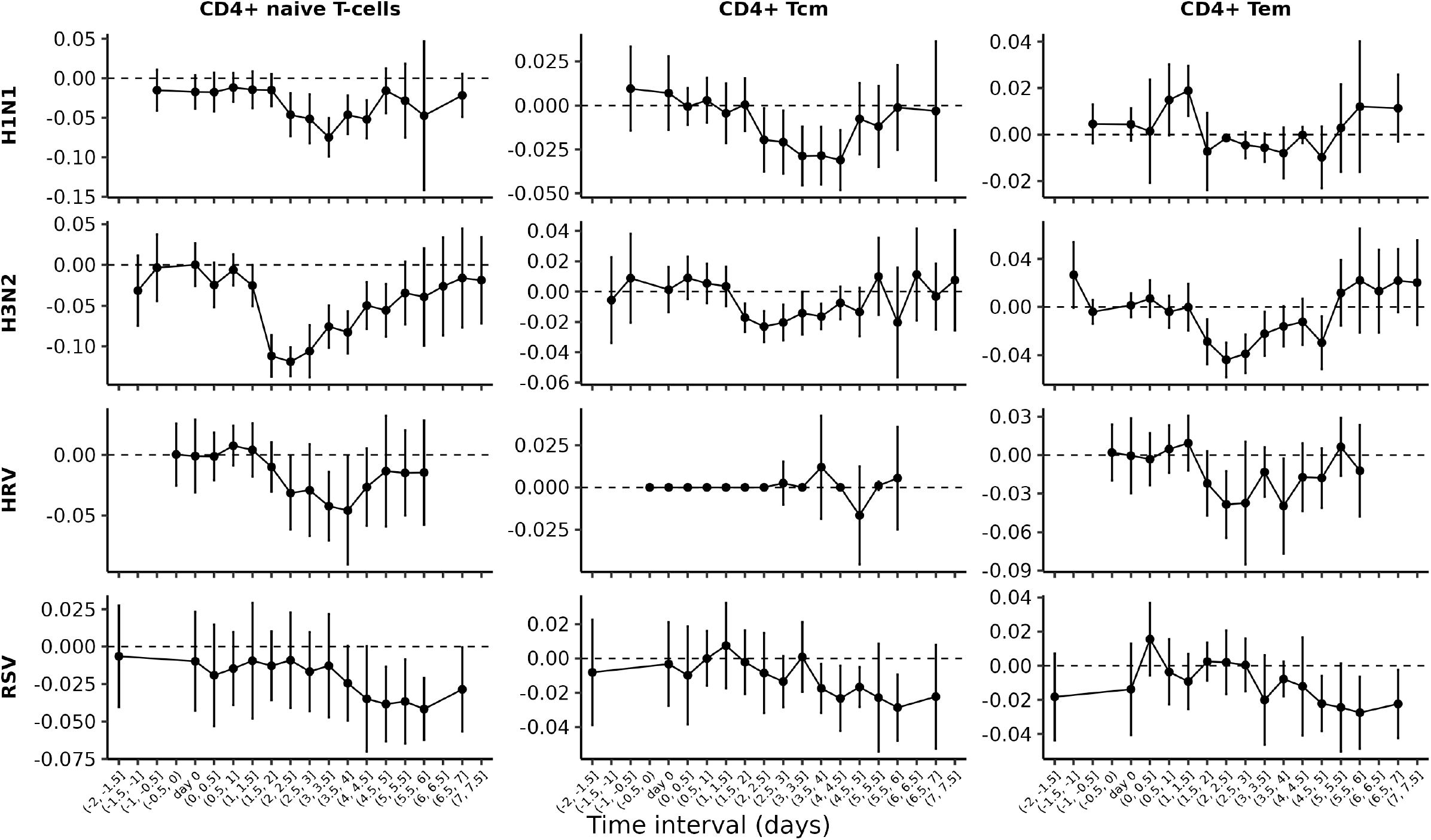
Fixed-effect meta-analysis of dendritic cell subtype enrichment following viral inoculation. Columns represent CD4+ naïve T cells (left), CD4+ central memory T cells (CD4+ Tcm, middle), and CD4+ effector memory T cells (CD4+ Tem, right). Rows correspond to viral pathogens, including H1N1, H3N2, HRV, and RSV (top to bottom). Each point shows the pooled effect size estimate from the fixed-effect meta-analysis comparing symptomatic/infected (Sx/Inf) versus asymptomatic/uninfected (Asx/Uninf) participants at each 0.5-day time interval. Vertical error bars denote 95% confidence intervals (CIs); intervals that do not cross zero indicate statistically significant differences between the two groups. A positive estimated value reflects higher relative enrichment in the Sx/Inf group. Day 0 represents the baseline time point prior to or at inoculation.

B-cell enrichment scores decreased significantly following challenge across all four viral studies, with the earliest reductions observed following H3N2 challenge and later responses observed following H1N1, HRV, and RSV challenge (Fig. 7). Similar temporal patterns were observed for class-switched memory B cells. In contrast, plasma-cell enrichment increased progressively following challenge across all four viral studies.

**Figure 7.**
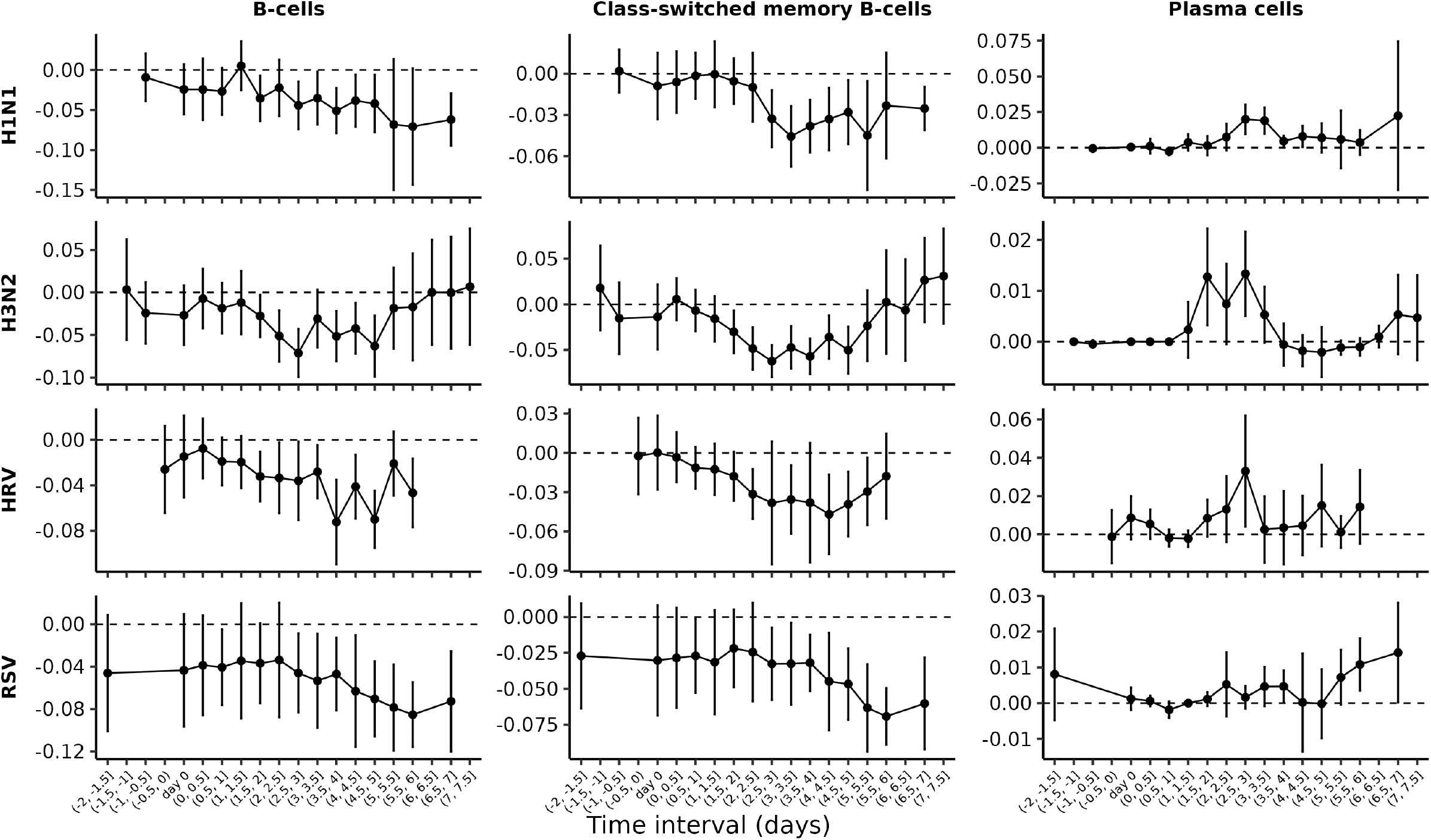
Fixed-effect meta-analysis of dendritic cell subtype enrichment following viral inoculation. Columns represent total B cells (left), class-switched memory B cells (middle), and plasma cells (right). Rows correspond to viral pathogens, including H1N1, H3N2, HRV, and RSV (top to bottom). Each point shows the pooled effect size estimate from the fixed-effect meta-analysis comparing symptomatic/infected (Sx/Inf) versus asymptomatic/uninfected (Asx/Uninf) participants at each 0.5-day time interval. Vertical error bars denote 95% confidence intervals (CIs); intervals that do not cross zero indicate statistically significant differences between the two groups. A positive estimated value reflects higher relative enrichment in the Sx/Inf group. Day 0 represents the baseline time point prior to or at inoculation.

Overall, these adaptive immune trajectories paralleled the earlier innate immune responses across studies, with H3N2 producing the most rapid immune shifts and RSV showing the slowest temporal kinetics.

### Virus-specific immune trajectories identify distinct respiratory viral response programs

Although many immune-cell trajectories were shared across respiratory viruses, several virus-specific patterns were identified. H3N2 challenge consistently produced the earliest and strongest immune-cell changes across both innate and adaptive compartments. Rapid increases in neutrophils, monocytes, and aDCs were detected before day 2 post-challenge, accompanied by early reductions in multiple T-cell and B-cell populations.

In contrast, RSV challenge showed delayed immune-cell dynamics across several cell populations. Significant increases in aDC enrichment were not observed until approximately day 6 post-challenge, and neutrophil enrichment increased later than in influenza studies. RSV studies also showed minimal pDC activation compared with H1N1 and H3N2 challenge studies. In the adaptive compartment, CD8^+^ Tcm enrichment remained consistently lower among Sx/Inf participants across much of the RSV study period.

HRV challenge exhibited additional distinct features. Neutrophil enrichment transiently decreased shortly after challenge, in contrast to the early neutrophil increases observed in H3N2 studies. HRV studies also showed weaker pDC responses and more modest changes in several lymphocyte populations compared with influenza challenge models.

Differences were also observed within adaptive immune-cell trajectories. Th1-cell enrichment increased significantly only following H3N2 challenge, whereas reductions in CD8^+^ Tem cells were less pronounced in RSV studies. Plasma-cell enrichment increased progressively across all viral studies, although the magnitude and timing varied between viruses.

Together, these findings indicate that respiratory viruses induce both shared and virus-specific immune trajectories, with differences observed in the timing, magnitude, and coordination of innate and adaptive immune responses. By aggregating results across multiple independent cohorts within the same viral challenge, the meta-analysis identified consistent temporal patterns across cohorts. **Migration-associated transcriptional programs reveal coordinated immune trafficking dynamics**

To investigate immune-cell populations associated with migration-related transcriptional programs, we examined the relationship between migration scores and xCell-derived cell-type enrichment scores using mixed-effects regression models (Fig. 8). In H3N2 studies, migration scores were positively associated with neutrophils, CD8^+^ Tem cells, and CD4^+^ T cells (FDR <0.05), with a marginal association observed for aDCs (FDR <0.1). In H1N1 studies, migration scores were significantly associated with neutrophils, aDCs, pDCs, CD4^+^ Tem cells, Th1 cells, and Th2 cells (FDR <0.05), whereas weaker associations were observed for CD8^+^ Tem cells.

**Figure 8.**
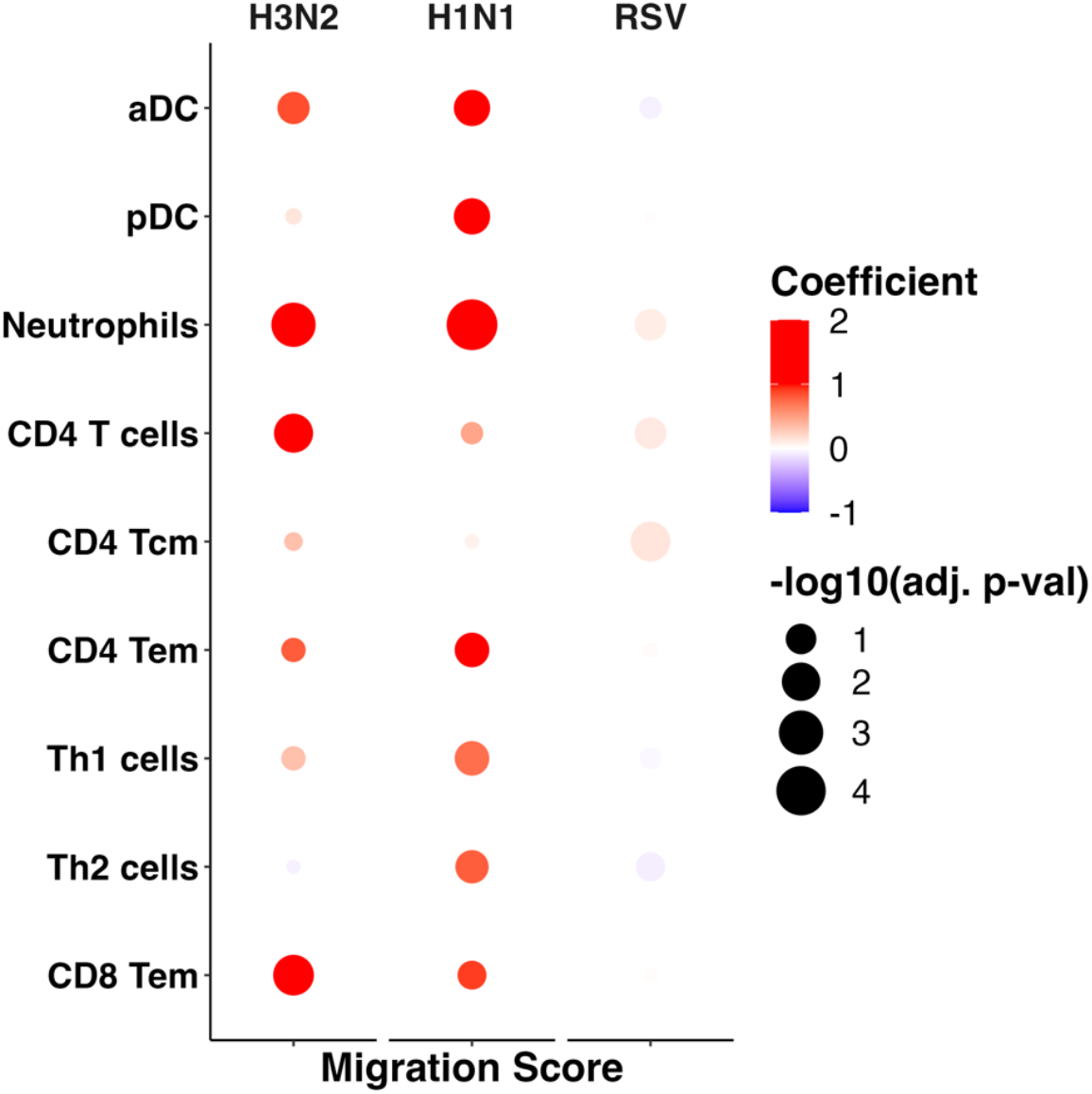
Association between migration score and cell type enrichment score. Mixed-effects models were run independently for the H3N2 (left), H1N1 (middle), and RSV (right) inoculation studies. Coefficients from the models are plotted, with higher positive coefficients indicated in red and lower negative coefficients in blue. The −log10(adjusted p-value) is represented by dot size, with smaller p-values corresponding to larger dots.

In RSV studies, migration scores showed a modest positive association with CD4^+^ Tcm enrichment, although the effect size was relatively small. Overall, both innate and adaptive immune-cell populations contributed to migration-associated transcriptional programs during respiratory viral challenges.

In contrast, residency scores were not significantly associated with enrichment of any cell type after multiple-testing correction in the H3N2, H1N1, or RSV studies. These findings suggest that dynamic migration-associated immune programs are more prominent than residency-associated programs in peripheral blood during acute respiratory viral infection.

## DISCUSSION

Using statistical deconvolution and xCell digital sorting of longitudinal bulk transcriptomic data, we examined immune-cell composition changes across human H3N2, H1N1, HRV, and RSV challenge studies. Despite differences between viruses, several broad patterns were consistently observed. Symptomatic infected participants generally showed early increases in innate immune-cell populations, particularly aDCs, pDCs, monocytes, and neutrophils, together with reductions in multiple T-cell and B-cell populations. However, the timing of these responses differed considerably between viruses. H3N2 challenge produced the earliest changes across many immune-cell populations, whereas RSV responses were delayed across both innate and adaptive compartments.

Baseline immune composition also differed between outcome groups before inoculation. Statistical deconvolution analyses showed that, compared with Asx/Uninf participants, Sx/Inf participants tended to exhibit higher neutrophil and monocyte proportions together with lower T-cell and B-cell proportions, particularly in the H3N2 studies. Consistent with these findings, xCell analyses identified similar baseline differences in several lymphocyte populations, including lower CD8^+^ T-cell and B-cell enrichment scores among Sx/Inf participants. Although not all of these differences reached statistical significance, the overall pattern suggests that pre-existing immune-state composition may contribute to susceptibility to symptomatic infection.

Trajectory analyses of innate immune populations showed that aDCs exhibited some of the earliest and strongest responses following influenza challenge. pDC enrichment also increased prominently following H1N1 and H3N2 challenge but remained relatively unchanged in RSV studies. Previous experimental studies have shown that RSV induces weaker type I interferon responses than influenza viruses and that pDC activation during RSV infection is comparatively limited (Jewell et al., 2007). Our findings in human challenge cohorts parallel these observations and further support delayed innate immune activation as a feature of RSV infection.

Although conserved immune trajectories were observed across H1N1, H3N2 and RSV challenges, HRV demonstrated several distinct immune features compared with influenza and RSV. In contrast to the broader and more coordinated innate activation observed following H1N1, H3N2, and RSV challenge, HRV responses were generally weaker or less synchronized across multiple innate immune populations, including pDCs, monocytes, neutrophils, and NK cells. These findings suggest that influenza viruses and RSV may share more common pathogen-induced systemic immune programs, whereas HRV induces a comparatively distinct host-response pattern.

Unlike innate immune cells, most T-cell and B-cell populations decreased following challenge, whereas plasma-cell enrichment progressively increased over time. One possible explanation is redistribution of circulating lymphocytes during acute infection, although other mechanisms, including apoptosis or altered trafficking patterns, may also contribute. Migration-associated transcriptional programs were observed across both innate and adaptive immune-cell populations, including neutrophils, aDCs, pDCs, CD8^+^ Tem cells, CD4^+^ Tem cells, Th1 cells, and Th2 cells, supporting active immune-cell trafficking during infection.

CD8^+^ Tem cells showed particularly notable differences between outcome groups in influenza studies. Previous work has shown that influenza-specific CD8 T-cell responses can exhibit substantial cross-reactivity across viral strains (Gras et al., 2010), and early CD8 effector responses have been associated with milder disease following natural influenza infection (Fox et al., 2012). Our analyses identified differences in CD8^+^ effector memory T-cell enrichment between outcome groups in influenza challenge studies, including at some pre-challenge time points, suggesting a potential role for pre-existing memory T-cell immunity in modulating infection outcomes.

We also observed selective increases in Th1-cell enrichment following H3N2 challenge. Mouse studies have shown that live influenza infection preferentially induces Th1-polarized responses, whereas inactivated influenza vaccination more commonly induces Th2-biased responses with weaker heterologous protection (Jung & Lee, 2020; Moran et al., 1999). Although vaccination responses were not examined here, the H3N2 findings are consistent with those observations.

An important aspect of this study is the integration of multiple independent human challenge cohorts using meta-analysis. By synthesizing results across independent cohorts for each virus, this approach enabled the identification of shared temporal immune patterns. Several limitations should be considered. First, all analyses were performed using peripheral blood transcriptomic data and may not fully reflect immune events occurring at respiratory mucosal sites. Second, some cohorts had relatively modest sample sizes, limiting power for smaller immune-cell differences. Finally, xCell enrichment scores and statistical deconvolution provide indirect estimates of cell composition and should ideally be validated using flow cytometry or single-cell approaches.

Overall, these analyses provide a comparative view of immune-cell dynamics across human respiratory viral challenge studies and highlight both shared and virus-specific patterns of immune activation across influenza viruses, HRV, and RSV.

