## Supplementary material for "Temporal High-Resolution Atlas of Human Blood Leukocyte Composition in Response to Respiratory Virus Inoculation": Supp. Table 1

| | | | |
|---|---|---|---|
| aDC | CD8+ naive T-cells | iDC | pDC |
| B-cells | CD8+ T-cells | Memory B-cells | Plasma cells |
| CD4+ memory T-cells | CD8+ Tcm | Monocytes | pro B-cells |
| CD4+ naive T-cells | CD8+ Tem | naive B-cells | Tgd cells |
| CD4+ T-cells | cDC | Neutrophils | Th1 cells |
| CD4+ Tcm | Class-switched memory B-cells | NK cells | Th2 cells |
| CD4+ Tem | DC | NKT | Tregs |

**Supplemental Table 1.** The 28 blood leukocyte cell subtypes out of referenced 64 cell types analyzed by xCell.
