## Supplementary material for "Temporal High-Resolution Atlas of Human Blood Leukocyte Composition in Response to Respiratory Virus Inoculation": Supp. Fig. 1

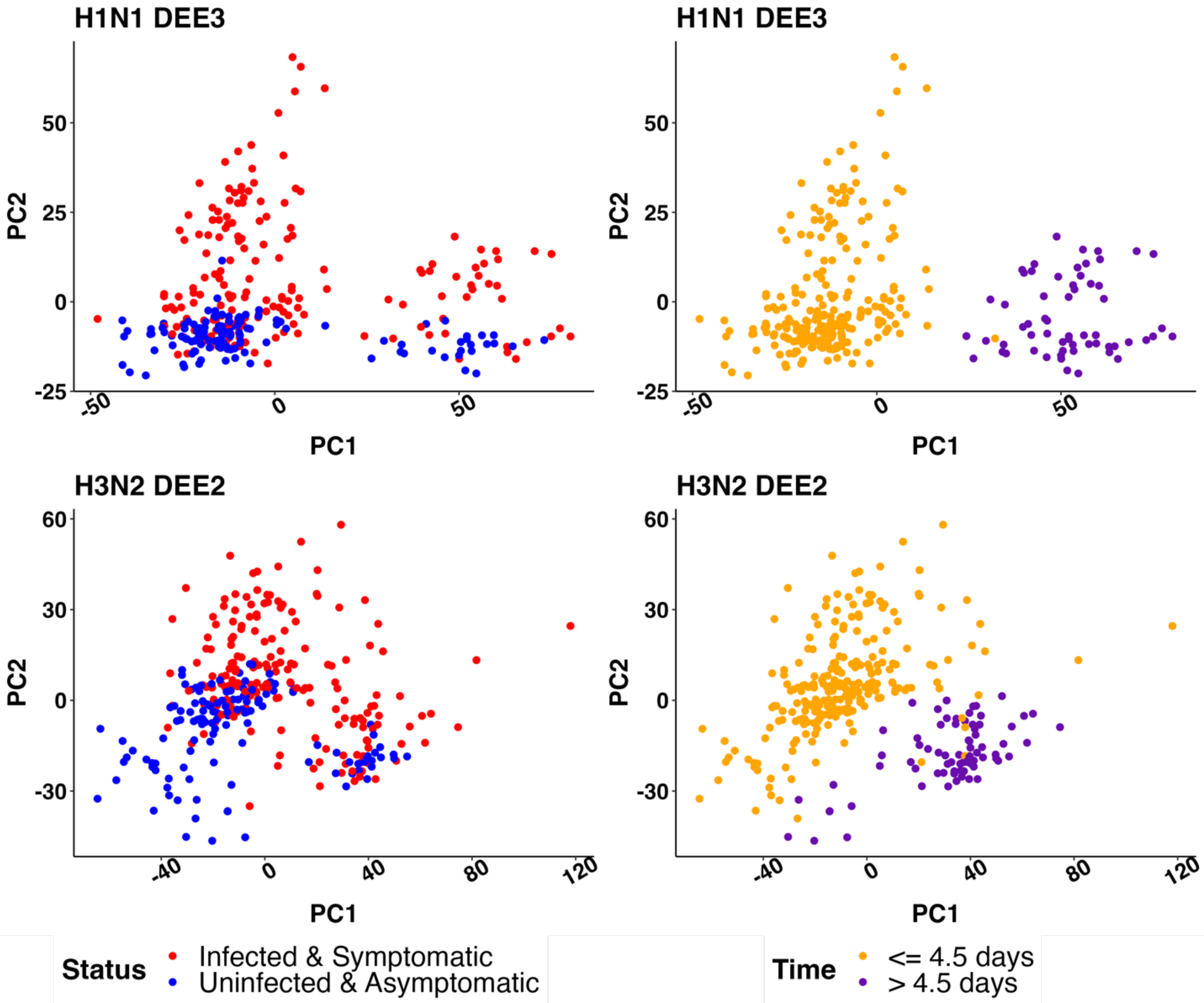


**Supplemental Figure 1**. Principal Component Analysis (PCA) on gene expression revealed distinct separations in one H1N1 (top) and one H3N2 (bottom) study from GSE73072. On the right, PCA shows clear separation by time, while the plots on the left depict principal components based on infection and symptom status.
